# simCRISPR: Modeling Experimental Complexity in Pooled CRISPR Screens

**DOI:** 10.64898/2026.05.14.725042

**Authors:** Zhaohan Zhu, Xiaoru Dong, Chanhee Kim, Christian Maugee, W. Brad Barbazuk, Christopher D. Vulpe, Rhonda Bacher

## Abstract

Pooled CRISPR screens are widely used to investigate gene function and uncover genetic interactions. However, benchmarking computational methods for detecting gene-by-environment (GxE) interactions remains difficult because ground truth is rarely available and existing simulation tools are not designed for GxE screening contexts. To address this, we developed `simCRISPR`, a flexible simulation framework for generating pooled CRISPR screen data under complex experimental designs. Using simulated datasets informed by empirical CRISPR screen designs, we evaluated commonly used analysis methods, comparing normalization strategies based on safe-harbor versus non-targeting sgRNAs and assessing empirical log_2_FC thresholds as an additional effect-size criterion. We found that safe-harbor-based normalization improved interaction detection when DNA damage-related effects were present, particularly when combined with empirical log_2_FC thresholding for DESeq2. Application of this workflow to a doxorubicin GxE screen further showed that safe-harbor-based normalization reduced bias in log_2_FC distributions and identified additional biologically relevant candidates. simCRISPR is available at https://github.com/bachergroup/simCRISPR.

## 1 Introduction

Pooled CRISPR screens enable systematic, genome-scale interrogation of gene function across diverse cell types and model systems. In a typical experiment, libraries of single-guide RNAs (sgRNAs) are delivered to cell populations to induce gene knockout or interference, and the resulting changes in cell survival or proliferation reveal genes essential for cellular processes. When paired with chemical or environmental exposures, CRISPR screens can also identify genes that modulate cellular responses to specific drugs or toxicants (Colic & Hart 2019). Here, we refer to such designs as gene-by-environment (GxE) CRISPR screens. These GxE interactions can reveal how genetic and environmental factors jointly shape cellular phenotypes.

Most existing analytical methods for CRISPR screens, however, were designed for simple essentiality studies that focus on gene-level main effects rather than interaction effects. In these designs, each gene is targeted by multiple sgRNAs across replicates in the absence of any additional chemical or environmental factors, and established analysis methods such as MAGeCK (Li et al. 2014, 2015) aggregate sgRNA-level count changes to estimate gene essentiality. Technical variability in sequencing depth across samples is accounted for by normalizing to non-targeting controls, which are sgRNAs designed not to match any genomic sequence in the target genome. In GxE screens, the same analysis framework is applied to identify conditionally essential genes by comparing knockout-only against knockout-plus-treatment samples, but it is unclear whether normalization to non-targeting controls remains valid in this setting.

More broadly, the choice of negative controls in pooled CRISPR screens has received increasing attention. Safe-harbor sgRNAs, which target transcriptionally inactive or phenotypically neutral genomic regions, have been proposed as alternative negative controls (Chen et al. 2018, Cheng et al. 2023, Morgens et al. 2017). The rationale is that guide binding and creating double strand breaks in the DNA may activate stress responses associated with DNA damage, thereby affecting cell survival and proliferation independent of the targeted gene. Since non-targeting guides do not capture this background signal, using them as negative controls can lead to improper normalization and false positives in downstream analysis (Chen et al. 2018, Cheng et al. 2023, Morgens et al. 2017).

In GxE screens, treatment-induced global responses, including DNA damage, stress signaling, or altered proliferation, may interact with both knockout effects and general guide binding effects. The performance of safe-harbor sgRNAs for normalization under these settings has not been systematically evaluated, partly because many datasets lack safe-harbor controls or include only a limited number, often less than a few dozen (Hayashi et al. 2020). At the same time, improved sgRNA design has increased knockout efficiency (Doench et al. 2016), allowing for fewer sgRNAs per gene. With less averaging across guides, gene-level estimates become more sensitive to global biases and therefore more dependent on robust normalization.

The combined effects of treatment-induced cellular responses, limited negative controls, and reduced sgRNAs per gene complicate normalization and analysis. Existing simulation frameworks, including CRISPulator (Nagy & Kampmann 2017), the benchmarking approach by Bodapati et al. (Bodapati et al. 2020), and gscreend (Imkeller et al. 2020) were developed primarily for simple essentiality screens and do not incorporate experimental features relevant to GxE screens. In particular, they do not support modeling treatment-by-knockout interaction effects and most do not model safe-harbor sgRNAs. As a result, their utility for benchmarking analysis methods or guiding experimental design is limited.

To better understand these challenges, we developed `simCRISPR`, a simulator for modern CRISPR screen designs that models treatment-by-knockout interactions, variability in sgRNA efficiency, both non-targeting and safe-harbor negative controls, and technical noise introduced by sample preparation and sequencing. We first compared `simCRISPR`-generated data with experimental datasets to demonstrate that our simulator captures key distributional characteristics observed in CRISPR screens. We then simulated multiple experimental scenarios to benchmark three widely used analysis methods and evaluated their performance under different normalization strategies. Based on these analyses, we provide practical recommendations for the design and analysis of pooled CRISPR screens aimed at detecting treatment-by-knockout interactions, including use of an empirical threshold to reduce false positives. `simCRISPR` is available as an R package for simulating pooled CRISPR screening data under various experimental designs. It can be downloaded via GitHub at https://github.com/bachergroup/simCRISPR.

## 2 Results

### 2.1 Overview of simCRISPR

In a typical GxE screen, cells are first genetically perturbed using an sgRNA library containing guides targeting genes for knockout and negative control guides. Some cells are then exposed to a chemical or environmental treatment, while others remain unexposed. After a period of proliferation, the two groups are sequenced as treatment (TRT) and control (KO) samples to evaluate resistance or sensitivity (Fig. 1A). To evaluate analysis methods under this experimental design, we developed `simCRISPR`, a simulator that incorporates both experimental and biological effects. The framework begins by simulating the underlying sgRNA distribution at baseline (T0), and then modeling the exponential growth process of the cell population according to sgRNA-specific growth parameters that are composed of genetic effects, exposure effects, genetic-by-exposure (GxE) interactions, DNA damage response (DDR) effects, and DDR-by-exposure (DDRxE) interactions. Following the growth model, cell counts are then subjected to technical variability introduced during sample preparation and sequencing (see Methods). The sgRNA-specific growth parameters for targeting, non-targeting, and safe-harbor guides are simulated based on the expected effects of each guide type in each sample type (Fig. 1B).

**Figure 1.**
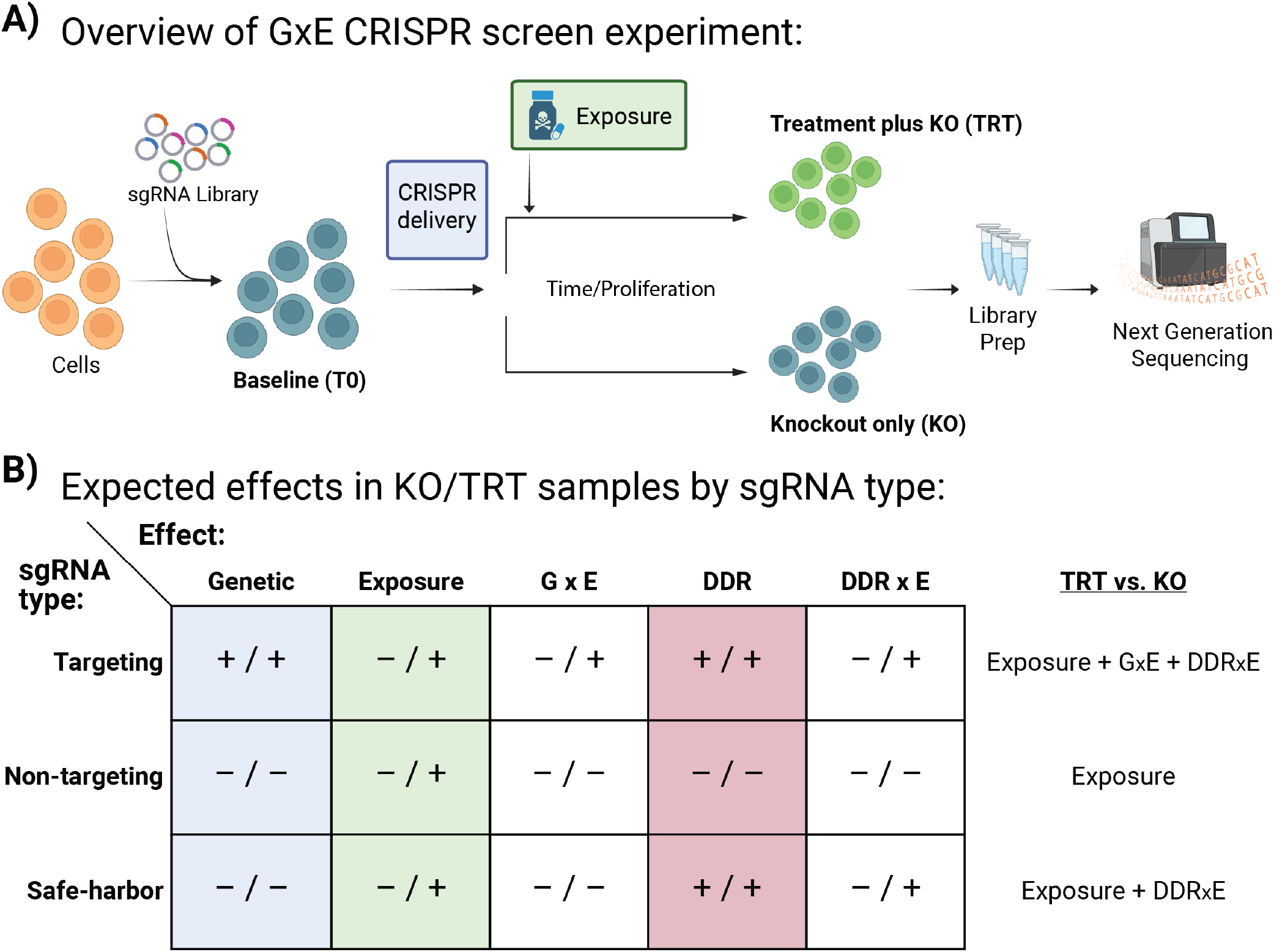
Overview of a GxE CRISPR screening experiment and expected effects. (A) Experimental design of a GxE CRISPR screen consisting of baseline (T0), knockout (KO), and treatment (TRT) sample types. (B) Presence (+) or absence (–) of effects in KO and TRT sample types by sgRNA type (formatted as KO / TRT). The TRT vs. KO column indicates which effect components remain when contrasting TRT with KO samples for each sgRNA type. Created in BioRender. Maugee, C. (2026) https://BioRender.com/wsa9a90.

Rather than providing an exact fit to a specific dataset, simCRISPR generates simulations under specified experimental designs using parameter values selected from systematically explored plausible ranges (see Methods). We compared simulated data to two empirical datasets, an essentiality screen and a GxE screen, to assess whether key distributional features were captured. The simple screen (HDCRISPR dataset) by Henkel et al., 2020 employed a genome-wide pooled sgRNA library containing an additional 300 non-targeting controls and 135 safe-harbor sgRNAs in HAP1-Cas9 cells (Henkel et al. 2020). Read counts were obtained from bulk HAP1-Cas9 cell populations and two single-cell-derived clones with high Cas9 editing activity. The GxE screen (Doxo dataset) was generated by Kim et al., 2026, in which HepG2/C3A cells underwent gene knockout perturbations under doxorubicin exposure to identify genetic modulators of chemical toxicity (Kim et al. 2026). One of the effects of doxorubicin specifically is activation of the DNA damage response pathway, which presents an interesting interaction to account for in downstream analysis.

Across the two different screens, raw read count distributions representing relative sgRNA abundance differed in overall mean and variability, consistent with differences in experimental design (Fig. 2A,B). Data simulated by `simCRISPR` under the Doxo design generally recapitulated sample count distributions (Fig. 2C). Baseline (T0) samples had the smallest variability, whereas KO samples showed wider distributions. TRT samples had slightly reduced variability compared to KO, possibly indicative of a selective pressure effect imposed under exposure.

**Figure 2.**
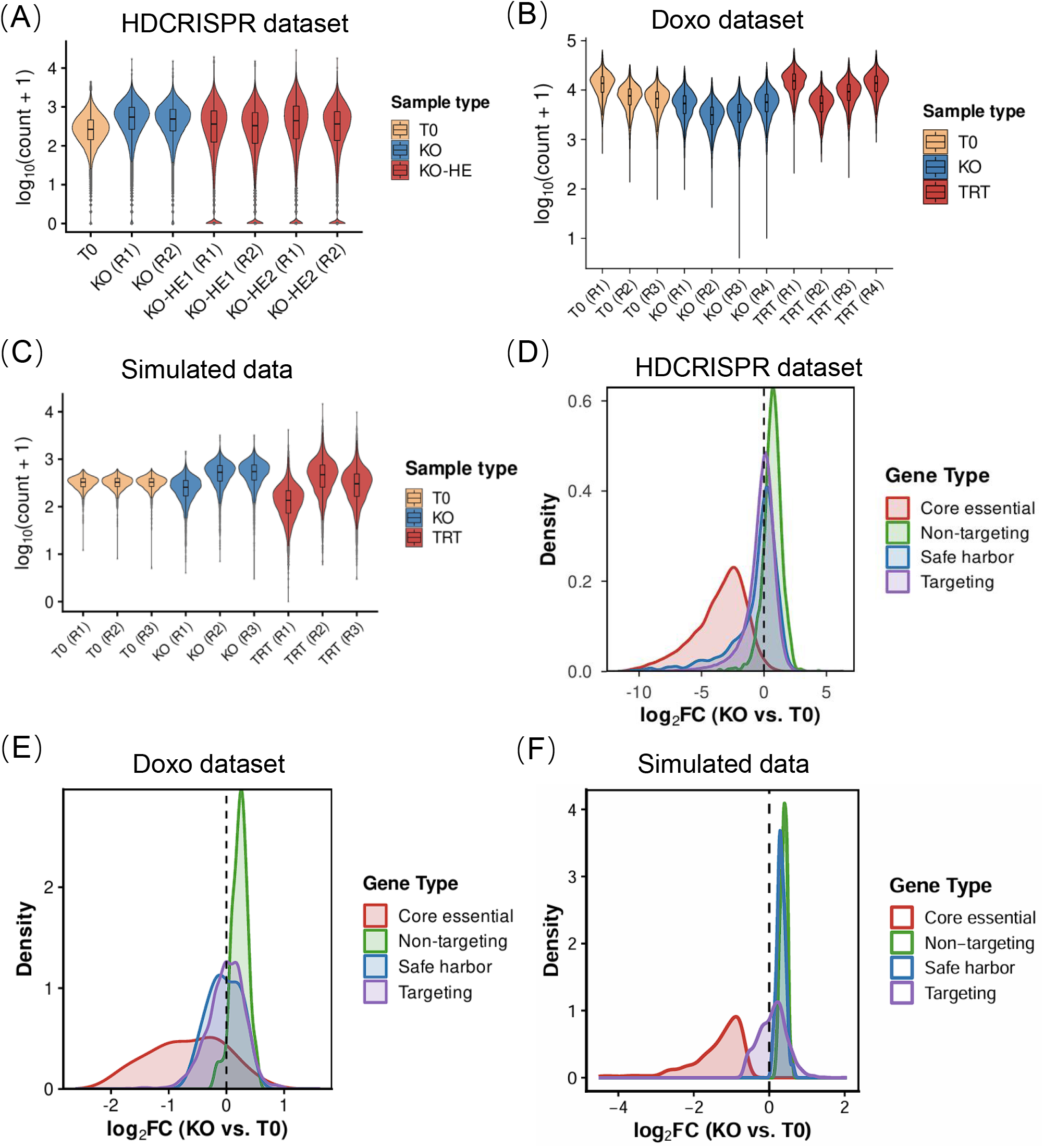
Comparison of empirical and simulated CRISPR screen data. (A) Violin plots of the raw read count distributions for each sample in the HDCRISPR dataset. (B) Violin plots of the raw read count distributions from the Doxo dataset. (C) Violin plots of simulated raw read counts from a representative simulation generated by `simCRISPR` based on the Doxo experimental design. (D) Density distributions of normalized log2 fold changes of sgRNA counts between KO and T0 across gene categories (core-essential, non-targeting, safe-harbor, and targeting) in the HDCRISPR dataset. (E) Density distributions of normalized log2 fold change for the corresponding gene categories in the Doxo dataset. (F) Density distributions of normalized log2 fold change for the corresponding gene categories in the representative simulated dataset.

We next examined distributions of the log_2_ fold-changes (log_2_FC) of normalized sgRNA counts between KO and T0 samples across sgRNA types. Using the original sgRNA category definitions as defined by Henkel et al. (2020), sgRNAs were grouped into four categories: non-targeting, safe-harbor, core-essential, and targeting. If non-targeting and safe-harbor sgRNAs were affected only by the exposure, both distributions should be centered around zero. In both screens, however, we observed separation between the two negative-control sgRNA groups, indicating the additional DDR-related effects in the safe-harbor guides (Fig. 2D,E). As expected, core-essential sgRNAs also showed a pronounced negative shift relative to negative controls, consistent with selection against essential gene loss. Reproducing these characteristics is therefore an essential feature of any credible simulation framework. Data simulated by `simCRISPR` recapitulated these log_2_FC distribution shifts across sgRNA types (Fig. 2F).

Having applied global normalization above, we next evaluated the effect of negative-control set choice on the targeting sgRNA log_2_FC distribution, as shown in Chen et al. (2018) (Chen et al. 2018). In the Doxo dataset, we compared targeting sgRNA log_2_FC values of sgRNA counts between TRT and KO samples under non-targeting and safe-harbor normalization (Fig. 3A). Non-targeting normalization produced a distribution centered slightly above zero, whereas safe-harbor normalization resulted in a clear leftward shift.

**Figure 3.**
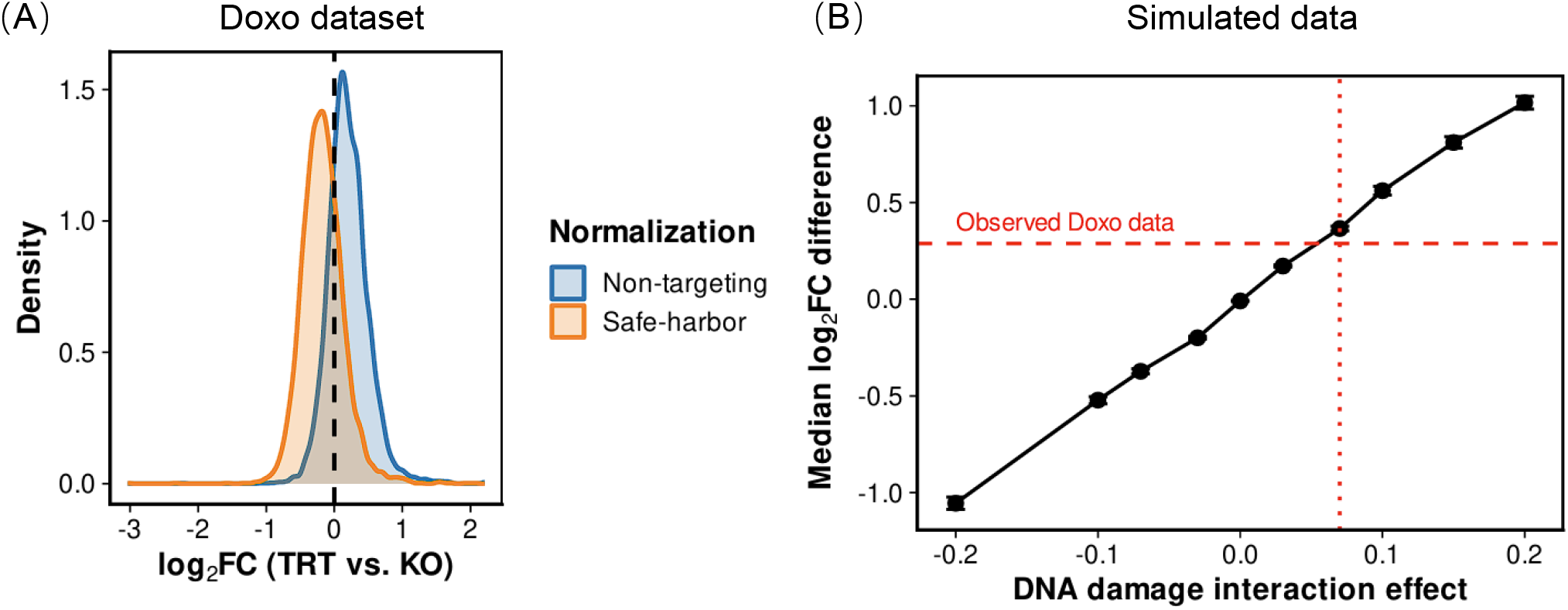
Normalization strategy affects log_2_FC distributions. (A) Distribution of normalized log_2_FC (TRT vs. KO) for targeting sgRNAs in the Doxo dataset under non-targeting and safe-harbor normalization. (B) Difference in median normalized log_2_FC (TRT vs. KO) between non-targeting and safe-harbor normalization across targeting sgRNAs is shown for different DNA damage-treatment interaction strengths. The dashed line shows the observed value in the Doxo dataset, and the dotted line indicates the corresponding interaction strength. Error bars represent variability across 25 simulation runs.

To assess whether this shift reflects DNA damage-treatment interaction (DDRxE), we simulated data with simCRISPR while varying the strength of the DDRxE effect. We captured the distributional shift by calculating the difference in median normalized log_2_FC between non-targeting and safe-harbor normalizations across all targeting sgRNAs. The median difference increased steadily with the simulated interaction effect (Fig. 3B), suggesting that stronger DNA damage-treatment interactions lead to greater separation between distributions obtained under the two normalization strategies.

### 2.2 Benchmarking interaction detection under different normalization strategies

Motivated by the differences in median log_2_FC (TRT vs. KO) observed between non-targeting (NT) and safe-harbor (SFHB) normalization, we next evaluated how normalization strategy affects GxE interaction detection across analysis methods. We considered two primary simulation settings, one with both DNA damage and DNA damage-treatment interaction effects, and one without, to establish baseline method performance across low, moderate, and high GxE effect sizes, where the GxE interaction parameter was set to be weaker than, comparable to, or stronger than the fixed environmental main effect parameter, respectively (see Supplementary File 1). We evaluated balanced accuracy in identifying conditionally essential sgRNAs across three widely used analysis strategies under either NT or SFHB normalization. To account for the practice of researchers combining statistical significance with effect-size filtering, we also evaluated a negative-control-based thresholded variant.

With no DDR or DDRxE effects, the balanced accuracy was similar between normalization strategies for each method across all levels of GxE effect (Fig. 4). Across methods, DESeq2 and MAGeCK-RRA showed comparable performance, whereas MAGeCK-MLE exhibited lower balanced accuracy at low GxE effect sizes but improved substantially at moderate and high GxE effect sizes. For DESeq2 and MAGeCK-RRA, balanced accuracy tended to decrease as the GxE effect size increased due to an increase in false positives among sgRNAs with small effects.

**Figure 4.**
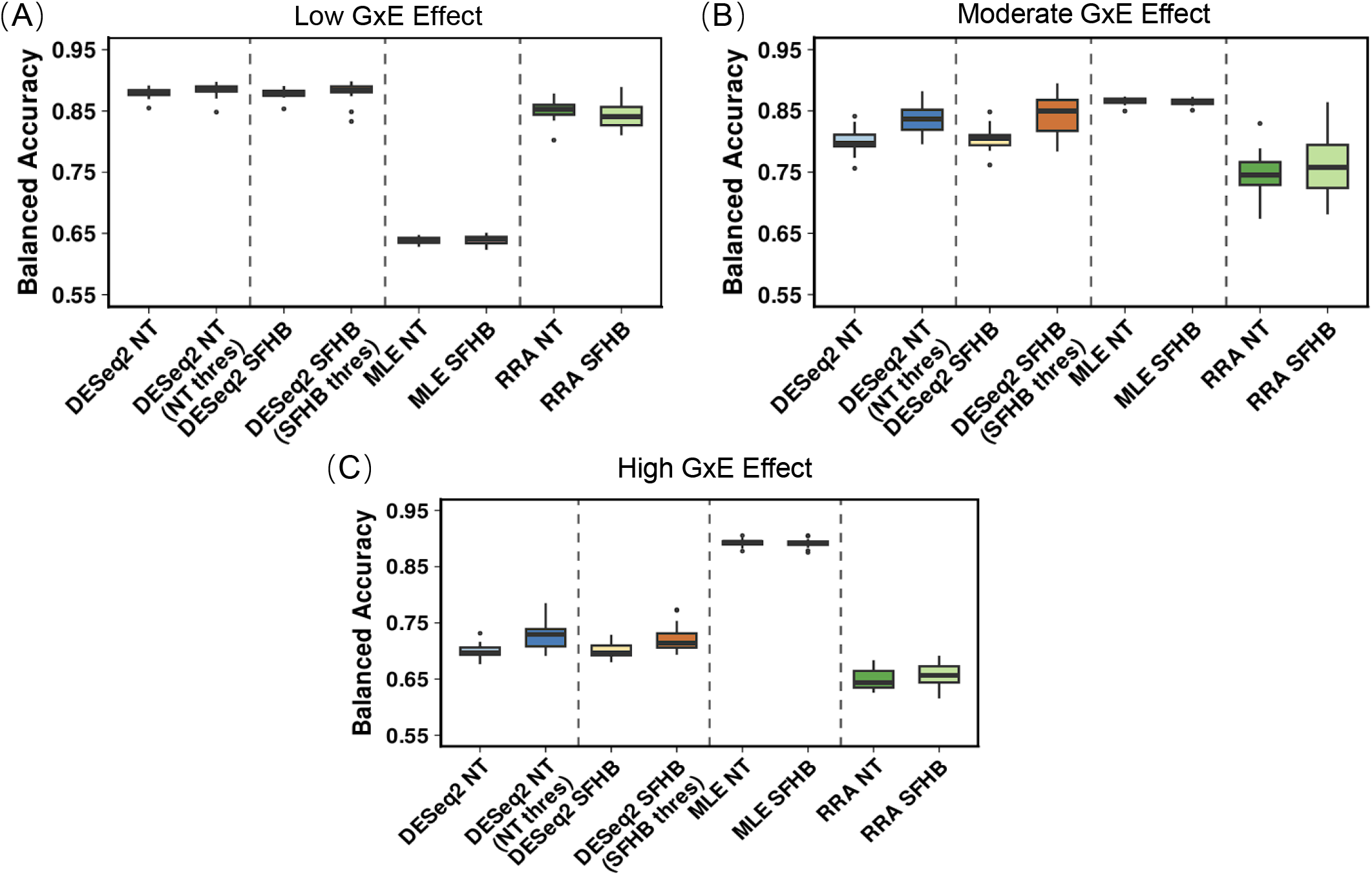
Method performance in the absence of DNA damage effects. Balanced accuracy of DESeq2, MAGeCK-MLE, and MAGeCK-RRA across low, moderate, and high GxE effect sizes under NT and SFHB normalization strategies. Empirical log_2_FC thresholding was applied only to DESeq2 results.

Using the same scenarios along with effects for DNA damage and DNA damage-treatment interactions, we observed that the choice of normalization strategy had a stronger influence on method performance (Fig. 5). SFHB-based normalization largely outperformed NT-based normalization across all GxE effect sizes. For MAGeCK-MLE, SFHB-based normalization yielded higher balanced accuracy at low and moderate GxE effect sizes, while the differences diminished at high effect sizes where strong interaction signals likely reduce the impact of normalization. For DESeq2, SFHB normalization showed a modest advantage over NT normalization at low GxE effect sizes, but this difference was not maintained at higher effect sizes, where performance was influenced by an increased number of false positives for sgRNAs with small effects. Applying an empirical log_2_FC threshold based on the negative control subsets substantially improved DESeq2 performance under SFHB normalization by removing candidates with small effects.

**Figure 5.**
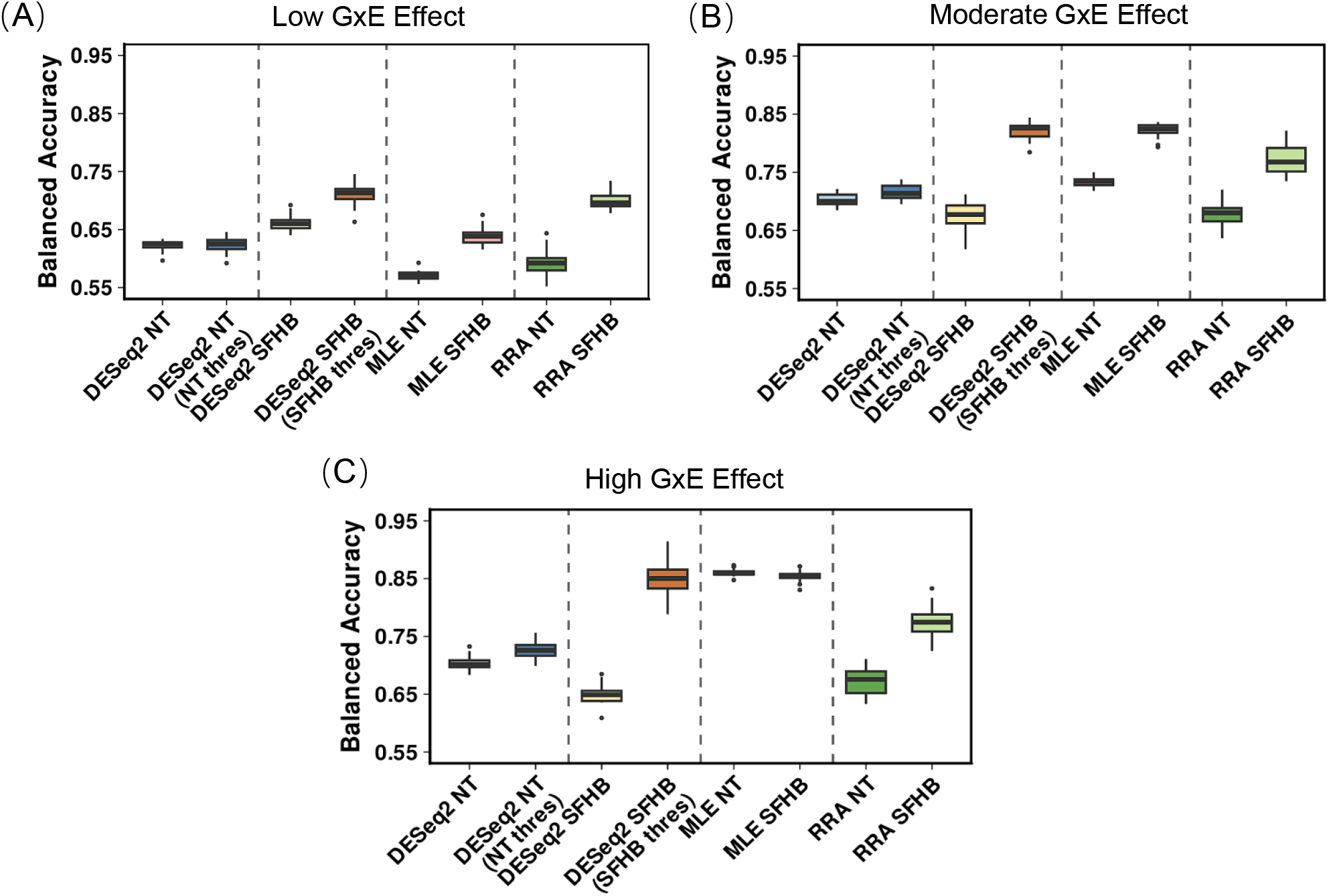
Method performance in the presence of DNA damage effects. Balanced accuracy of DESeq2, MAGeCK-MLE, and MAGeCK-RRA across low, moderate, and high GxE effect sizes under NT and SFHB normalization in simulations with DNA damage and DNA damage-treatment interaction effects. Empirical log_2_FC thresholding was applied only to DESeq2 results.

### 2.3 Application to DNA damage response (DDR) empirical data

Given the consistently strong performance of DESeq2 with SFHB-based normalization and thresholding in the presence of non-exposure-related DDR effects, we proceeded to use this approach to analyze the Doxo dataset (Kim et al. 2026) (Fig. 6). For comparison, we contrasted this approach with a practical scenario in which safe-harbors are unavailable, using DESeq2 with NT-based normalization and thresholding.

**Figure 6.**
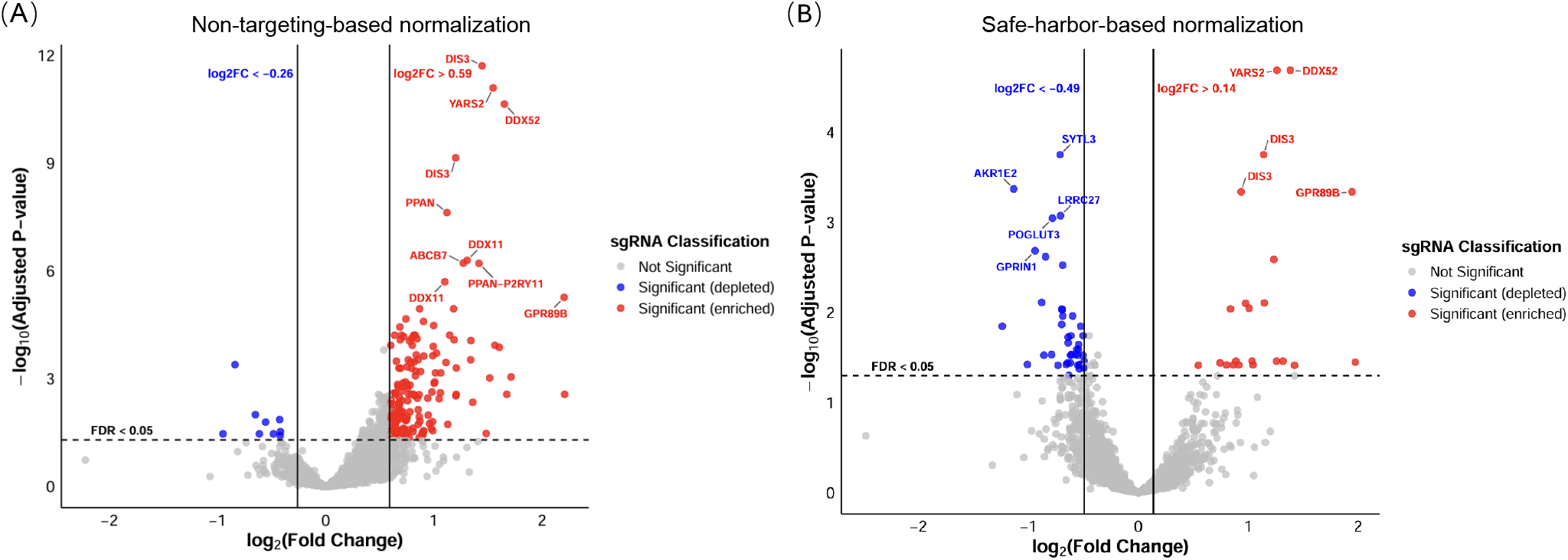
Application to Doxo GxE CRISPR screening dataset. **(A)** Volcano plot based on DESeq2 analysis under NT normalization and thresholding. **(B)** Volcano plot based on DESeq2 analysis under SFHB normalization and thresholding. Red and blue indicate significantly enriched and depleted sgRNAs, respectively, and gray points denote non-significant sgRNAs.

With the NT-based approach, log_2_FC (TRT vs. KO) values across all 3,206 sgRNAs were strongly right-shifted: 2,445 sgRNAs (76.2%) showed positive log_2_FC values, whereas only 761 sgRNAs (23.7%) showed negative log_2_FC values (Fig. 6A). Consistent with this global imbalance, the significant sgRNAs exhibited a pronounced right-skew, with 181 sgRNAs (5.6% of all sgRNAs; 95.3% of significant sgRNAs) showing significantly positive log_2_FC values, compared to only 9 sgRNAs (0.3% of all sgRNAs; 4.7% of significant sgRNAs) showing significantly negative log_2_FC values (Fig. 6A). In contrast, under the SFHB-based approach, which we prioritized based on its consistently stronger simulation performance, the log_2_FC values across all sgRNAs were more evenly distributed, with 2,104 sgRNAs (65.6%) showing negative log_2_FC values and 1,102 sgRNAs (34.4%) showing positive log_2_FC values. This improved balance was also reflected among significant sgRNAs, where 41 sgRNAs (65.1% of significant sgRNAs) were depleted and 22 sgRNAs (34.9% of significant sgRNAs) were enriched, resulting in a more symmetric distribution around zero (Fig. 6B).

Under SFHB normalization, 31 additional depleted sgRNAs were identified compared to NT normalization. One of these, CBR3 (carbonyl reductase 3), has been shown in prior studies to be directly linked to doxorubicin metabolism and doxorubicin-induced toxicity (Fan et al. 2008).

## 3 Discussion

In this study, we developed `simCRISPR`, a simulation framework for modern pooled CRISPR screens that models knockout effects, treatment effects, and their interactions, while incorporating key sources of technical variability such as DNA damage responses, PCR amplification bias, and sequencing noise. By benchmarking commonly used analysis methods under varying simulation settings, we provide both a practical tool for method evaluation and a principled framework for examining how experimental design and normalization choices influence the detection of interaction effects.

We used `simCRISPR` to evaluate the impact of normalization strategy under different simulation settings. In the absence of DNA damage and DNA damage-treatment interaction effects, NT- and SFHB-based normalization showed similar performance across methods, suggesting that either approach is sufficient when global editing-related effects are minimal. In contrast, when DNA damage and DNA damage-treatment interaction effects were introduced, SFHB-based normalization improved performance across DESeq2, MAGeCK-RRA, and MAGeCK-MLE, indicating that non-targeting sgRNAs do not fully capture DNA editing-related background effects. Although non-targeting sgRNAs are widely used in practice, our results support the routine inclusion of safe-harbor sgRNAs to provide more stable normalization and more reliable GxE effect estimation in CRISPR screens.

Based on our evaluations on simulated scenarios, we identified DESeq2 with SFHB-based normalization followed by SFHB-based thresholding as the preferred strategy, as it performed best across a range of GxE effect sizes. We applied this workflow to the Doxo dataset and compared it to using DESeq2 assuming no safe-harbors are available, reflecting a practical analysis scenario (i.e., NT-based normalization and thresholding). The SFHB-based approach better recovered established DDR-related genes and identified additional candidate genes that were not highlighted by the NT-based approach. These results indicate that the proposed workflow improves biological interpretability by reducing the impact of global treatment effects and enables more reliable identification of candidate genes.

We do note some limitations of our study. First, although `simCRISPR` incorporates a wide range of biological and technical effects, simulation outcomes necessarily depend on the choice of parameter settings. While we explored a broad range of scenarios and effect sizes, future extensions of the framework could explore dataset specific parameter estimation procedures. Additionally, safe-harbor sgRNAs are not included in all CRISPR libraries; additional work in defining appropriate negative controls, such as sgRNAs with stable behavior across conditions may be needed to support downstream analyses in such cases.

## 4 Methods

### 4.1 Overview

We designed a simulation framework that recapitulates the CRISPR experimental process for GxE screens. The simulation process assumes the dataset consists of *p* samples and *n* single-guide RNAs (sgRNAs), with the count *y*_*ij*_ representing the relative abundance of sgRNA *i* in sample *j*, where 1 ≤ *i* ≤ *n* and 1 ≤ *j* ≤ *p*. Each sample belongs to either the baseline (T0), control (KO only) group (*j* ∈ *J*_ctrl_) or the treatment (KO plus exposure) group (*j* ∈ *J*_trt_). We assume the sgRNA library includes up to three sgRNA categories: non-targeting sgRNAs (*i* ∈ *I*_ntgt_), safe-harbor sgRNAs (*i* ∈ *I*_sfhb_), and targeted knockout sgRNAs (*i* ∈ *I*_tgt_).

Non-targeting sgRNAs primarily capture technical variation and global exposure-associated effects, while safe-harbor sgRNAs additionally account for general editing-related effects such as Cas9 activity and potential DNA damage response (DDR). Targeted knockout sgRNAs target genes with the intention of perturbing cellular processes, potentially altering growth dynamics or other measurable phenotypes.

In our simulation framework, we focus on cell proliferation/survival outcomes, which are most commonly measured in CRISPR screens. We simulate data by modeling sgRNA-specific proliferation effects based on sgRNA categories and sample-specific expected effects. The simulated sample counts are also subjected to technical effects in sample preparation and sequencing.

#### 4.1.1 Simulation model

We first assume that sgRNA abundance follows an exponential growth process (Imkeller et al. 2020). Specifically, the count for sgRNA *i* in sample *j* at time *t* may be modeled as:

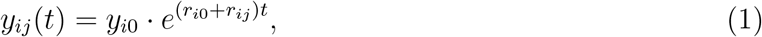

where *y*_*i*0_ ∼ N (*µ*^0^, (*σ*^0^)^2^) denotes the initial abundance of sgRNA *i*, with *µ*^0^ representing the mean initial sgRNA abundance and *σ*^0^ representing the standard deviation of initial sgRNA abundance across the library. The term *r*_*i*0_ is the intrinsic growth rate of cells with sgRNA *i*, capturing general sgRNA-specific growth effects such as efficiency shared across all samples. *r*_*ij*_ represents a sample- and sgRNA-specific deviation in growth that accounts for the experimental design and sgRNA type. *r*_*ij*_ consists of five effect components:

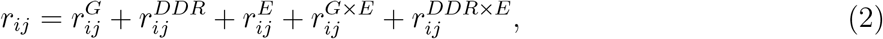

Where 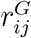 denotes the genetic perturbation effect, 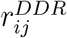 captures the DNA damage response, 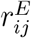 represents the exposure effect, 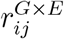 corresponds to the gene-by-exposure interaction, and 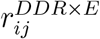 models the interaction between DNA damage response and exposure. This decomposition is, by design, adaptable to simulate data under alternative designs and assumptions.

#### 4.1.2 Parameter generation

In observed CRISPR screens, the intermediate values and processes are not observable, thus directly estimating parameter values is challenging. Instead, we focused on reflecting the underlying data generation process with flexible hyper-parameters to simulate a wide range of assumptions.

The genetic perturbation effect 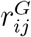 is generated using a mixture of signed exponential distributions to reflect that some genetic perturbations enhance proliferation while others inhibit it. Specifically, each sgRNA is first assigned to one of two effect directions representing growth-enhancing or growth-inhibiting perturbations. Let 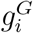 ∼ Bernoulli(*π*^*G*^), where 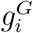 ∈ {0, 1} such that 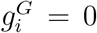 corresponds to growth-inhibiting effects and 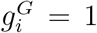 corresponds to growth-enhancing effects. Parameter *π*^*G*^ controls the proportion of sgRNAs with growth-enhancing effects.

Conditional on the direction assignment, the magnitude of the genetic perturbation effect (denoted by 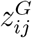 ) is drawn from an exponential distribution: 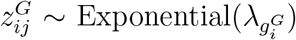, where the rate parameter 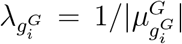 determines the expected magnitude of the effect for group 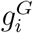 . Here, 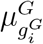 is the group-specific genetic perturbation effect parameter, with 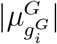 corresponding to the expected magnitude of the exponential distribution. The genetic perturbation effect is then defined as

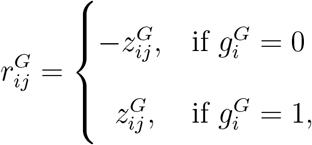

This formulation captures asymmetric and heavy-tailed effect size distributions commonly observed in CRISPR screens.

For non-targeting and safe-harbor sgRNAs (*i* ∈ *I*_ntgt_ ∪ *I*_sfhb_), the genetic perturbation effect is fixed at zero, i.e., 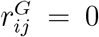, assuming these sgRNAs are not associated with gene perturbation effects.

The DNA damage response effect 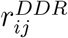 reflects mild cellular responses, such as those triggered by Cas9-induced double-strand breaks. For sgRNAs in the target or safe-harbor groups (*i* ∈ *I*_tgt_ ∪ *I*_sfhb_), we sample the raw DNA damage effect as:

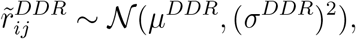

where *µ*^*DDR*^ and *σ*^*DDR*^ are user-specified simulation parameters that control the mean and variability of the raw DNA damage response effect, respectively. The final DNA damage response effect is then defined as

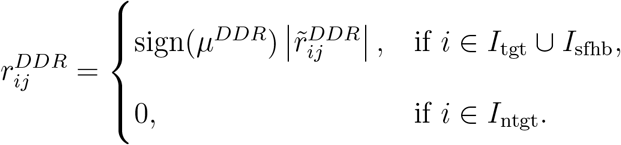

This formulation constrains the DNA damage response to have a consistent direction determined by the sign of *µ*^*DDR*^. In practice, because the DNA damage response is modeled as a global background effect, *σ*^*DDR*^ is typically set to be small, allowing only modest variation in effect magnitude across affected sgRNAs.

The exposure effect 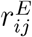 is applied across all sgRNAs in TRT samples. To model a consistent directional exposure effect, we sample

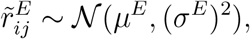

and then define

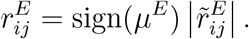

Thus, the direction of the exposure effect is determined by the sign of *µ*^*E*^; for example, a negative *µ*^*E*^ represents a globally inhibitory exposure effect. The exposure effect is set to zero in non-TRT samples.

The gene-by-exposure interaction effect 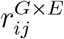 is modeled for target sgRNAs in TRT samples. For *i* ∈ *I*_tgt_ and *j* ∈ *J*_trt_, each sgRNA *i* is first assigned to one of two interaction groups. Let

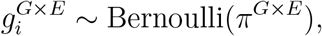

where 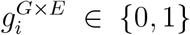 indicates the direction membership of sgRNA *i*, and *π*^*G×E*^ controls the proportion of sgRNAs with exposure-resistant interaction effects.

Conditional on the direction assignment, the magnitude of the interaction effect (denoted by 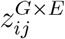) is sampled from an exponential distribution:

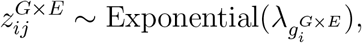

where 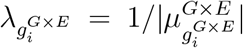 determines the expected magnitude of the interaction effect. Here, 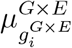 is the direction-specific interaction effect parameter, with 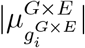 corresponding to the expected magnitude of the exponential distribution. The interaction effect is then defined as

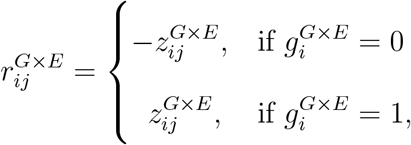

where 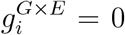 corresponds to exposure-sensitive interaction effects and 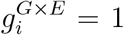 corresponds to exposure-resistant interaction effects. For all other sgRNAs and non-TRT samples, the GxE interaction effect is set to zero.

Similarly, the DDR-by-exposure interaction effect 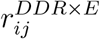 is included for sgRNAs in the target or safe-harbor groups in TRT samples. For *i* ∈ *I*_tgt_ ∪ *I*_sfhb_ and *j* ∈ *J*_trt_, each sgRNA *i* is first assigned to one of two interaction directions. Let

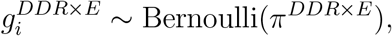

where 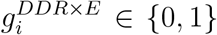 indicates the direction membership of sgRNA *i*, with 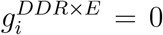 corresponding to DDR-by-exposure sensitive effects and 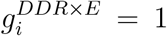 corresponding to DDR-by-exposure resistant effects. The parameter *π*^*DDR×E*^ controls the proportion of sgRNAs assigned to the DDR-by-exposure resistant direction.

Conditional on the direction assignment, the magnitude of the interaction effect (denoted by 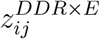) is sampled from an exponential distribution:

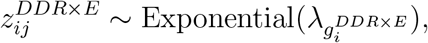

where 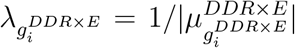 determines the expected magnitude of the interaction effect. The interaction effect is then defined as:

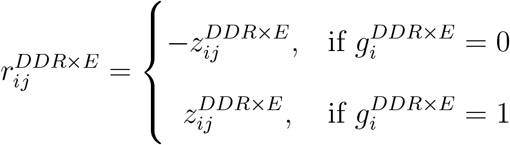

where 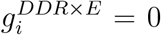 and 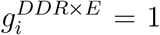 correspond to the two DDR-by-exposure interaction directions. In practice, *π*^*DDR×E*^ can be set to one, resulting in a consistent direction for the DDR-by-exposure interaction effect, as used in our simulation settings. For all other sgRNA and sample combinations, the DDR-by-exposure interaction effect is set to zero.

Since the distributional parameters of the genetic perturbation effects cannot be directly inferred from observed data, we generated simulated data under a wide variety of values to examine the performance of downstream analysis methods (see Supplementary File 2).

#### 4.1.3 Sequencing effect

In addition to biological effects, CRISPR screens undergo additional sample processing that may introduce technical variability in the observed counts between samples. We additionally capture these in our simulation framework. During sequencing library preparation, sgRNAs undergo amplification using polymerase chain reaction (PCR). To model this step, we define a PCR efficiency matrix *E* ∈ ℝ^*n×p*^, where each entry *E*_*ij*_ represents the amplification efficiency of sgRNA *i* in sample *j*. For each sample *j*, the PCR efficiency vector *E*_*·j*_ is sampled from a multivariate normal distribution:

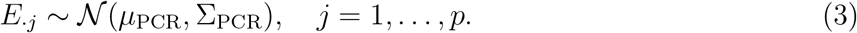

where Σ_PCR_ is an *n* × *n* diagonal covariance matrix with diagonal entries 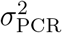 The mean vector *µ*_PCR_ has entries drawn from 𝒩 (*µ*_*cm*_, (*σ*_*cm*_)^2^). Following *r* rounds of PCR, the amplified fragment count *A*_*ij*_ is calculated as

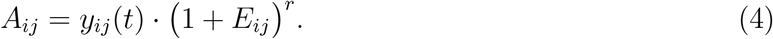

Next, we modeled the sequencing process by incorporating technical variation in library pooling. To mimic experimental equalization of input material, each sample is rescaled to match the smallest library size across samples. Let *M*_*j*_ = Σ_*i*_ *A*_*ij*_ denote the total amplified fragments in sample *j*, and a scaling factor is computed as

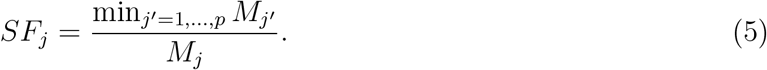

To reflect variance in this procedure, such as pipetting error or imprecise mixing, we introduce a small amount of Gaussian noise 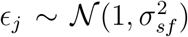 to form 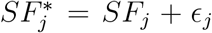 . The adjusted library size is 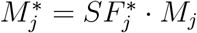. The equalized fragment counts for sample *j*, denoted 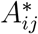, are sampled as

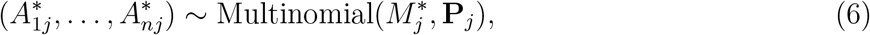

where **P**_*j*_ = (*P*_1*j*_, …, *P*_*nj*_) represents each sample’s sgRNA relative abundances:

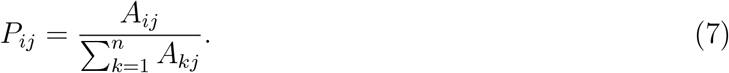

After equalization, the libraries are pooled into a single sequencing library. The sequencing probability for sgRNA *i* in sample *j* is defined as

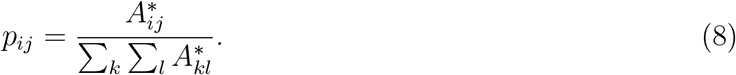

Finally, the observed read counts are generated by sampling from a multinomial distribution with total sequencing depth *R*. The observed count for each sgRNA *i* in sample *j*, denoted *C*_*ij*_, is derived from

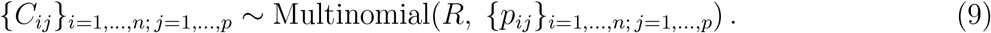

### 4.2 Negative controls and normalization strategy

Normalization is required to account for differences in sequencing depth and global shifts in sgRNA abundance across samples. Following the DESeq2 framework (Love et al. 2014), sample-wide scaling factors were computed using a set of negative control sgRNAs in all analyses with the exception of the data shown for Figure 2 (where scaling factors were computed on the entire set of sgRNAs).

### 4.3 Differential analysis methods

To evaluate the interaction effects between gene knockout and treatment conditions, we evaluated three established statistical frameworks for pooled CRISPR screens: DESeq2 (Love et al. 2014), MAGeCK RRA (Li et al. 2014), and MAGeCK MLE (Li et al. 2015).

#### DESeq2

We analyzed differential sgRNA abundance using the *DESeq2* package (v1.42.0) implemented in R (v4.3.2), which models count data under a negative binomial framework to account for biological and technical variation across replicates. Simulated count matrices were imported with the *DESeqDataSetFromMatrix()* function, specifying the design formula *∼condition* to contrast KO and TRT samples. Size factors, which are included as offsets in DESeq2’s model, were estimated using the *estimateSizeFactors()* function, with either non-targeting or safe-harbor sgRNAs designated as negative controls. Differential testing was performed through the *DESeq()* function, and resulting statistics were extracted with the *results()* function. *p*-values were adjusted using the Benjamini-Hochberg procedure to control the false discovery rate, yielding sgRNA-level responses specific to gene-treatment interactions.

#### MAGeCK RRA

We used the *MAGeCK RRA* package (v0.5.9.4) implemented in Python to identify sgRNA-level enrichment or depletion under treatment exposure (Li et al. 2014). Analyses were performed using the *mageck test* command, with treatment and control replicates specified by the `-t` and `-c` options, respectively. Normalization was performed using the `--norm-method control` parameter, where either non-targeting or safe-harbor sgRNAs were provided as negative controls (`--control-sgrna`). To minimize bias from uninformative guides, zero-count sgRNAs were excluded when present using `--remove-zero both` and `--remove-zero-threshold 0`. MAGeCK applies a robust rank aggregation (RRA) framework to summarize sgRNA-level evidence across replicates and generate significance statistics for downstream analysis. Resulting *p*-values were adjusted using the Benjamini-Hochberg procedure to control the false discovery rate. For downstream evaluation, sgRNA-level summary statistics were extracted from the MAGeCK output.

#### MAGeCK MLE

We applied the *MAGeCK MLE* package (v0.5.9.4), implemented in Python to estimate sgRNA-level coefficients across multiple experimental conditions (Li et al. 2015). This framework fits a generalized linear model under a negative binomial assumption to estimate coefficients corresponding to the baseline, knockout, and treatment conditions from sgRNA count data. Analyses were performed using the *mageck mle* command, with the count matrix (`-k`), design matrix (`-d`), and output prefix (`-n`) specified for each simulation run. Normalization was conducted using non-targeting or safe-harbor sgRNAs as negative controls (`--control-sgrna`), through control-based scaling (`--norm-method control`) to correct for sequencing depth and global treatment shifts. Model coefficients (*β* values) were estimated using ten computational threads (`--threads 10`) and two permutation rounds (`--permutation-round 2`) to ensure numerical stability. Design matrices included a baseline column and condition indicator columns for KO and TRT samples, with T0 samples defining the baseline reference condition. Thus, *β*_KO_ and *β*_TRT_ represent the estimated KO-versus-T0 and TRT-versus-T0 coefficients, respectively. The GxE interaction effect was calculated as the TRT-versus-KO contrast, *β*_TRT_ − *β*_KO_, and used for downstream evaluation.

### 4.4 Definition of true interaction labels

In the simulation framework, interaction effects were generated as continuous signed effect sizes. For performance evaluation, we converted them into ground-truth interaction labels using a threshold based on the simulated interaction effect magnitude. In the baseline analyses, this threshold was defined as

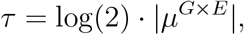

where *µ*^*G×E*^ denotes the GxE interaction-effect parameter used in the corresponding simulation setting. Because the interaction effect magnitude followed an exponential distribution with rate 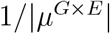, this threshold corresponds to the median magnitude of the simulated interaction effect under that setting.

An sgRNA was defined as a true interaction hit if 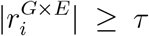, and as background otherwise. These binary labels were used to compute balanced accuracy.

### 4.5 Evaluation metrics

To compare the performance of different statistical methods under various normalization strategies, we used balanced accuracy (BA). Balanced accuracy is defined as the mean of sensitivity and specificity,

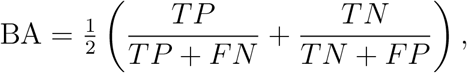

which accounts for class imbalance between sgRNAs with and without true interaction effects (Brodersen et al. 2010). Balanced accuracy was computed from confusion matrices constructed based on the simulated ground truth interaction labels.

### 4.6 Empirical cutoff estimation from negative controls

After normalization using negative control sgRNAs, sgRNA-level effect size estimates (log_2_ fold changes) and Benjamini-Hochberg adjusted p-values (FDR) were obtained from the corresponding statistical models. Statistical significance was initially assessed using a criterion of FDR *<* 0.05. To further incorporate effect-size information in a data-driven manner, we derived empirical log_2_ fold-change thresholds using negative control sgRNAs to approximate the background distribution of log_2_FC values. Negative control sgRNAs (non-targeting or safe-harbor sgRNAs) were first stratified by effect direction (log_2_FC *<* 0 or log_2_FC *>* 0). Within each stratum, we calculated the 2.5th percentile of the FDR distribution and used these values to define an empirical null set by retaining sgRNAs with FDR values greater than or equal to the corresponding directional cutoff. Using this null set, we then estimated lower and upper effect-size thresholds as the 2.5th and 97.5th percentiles of log_2_ FC, respectively, corresponding to the central 95% interval of background variation. Final candidate sgRNAs were required to satisfy both statistical significance (FDR *<* 0.05) and an effect-size criterion exceeding the empirical log_2_FC thresholds. This combined criterion ensures that selected sgRNAs are not only statistically significant but also exceed empirically defined effect-size thresholds.

### 4.7 Data sources and parameter settings

The full set of simulation parameters used to generate datasets resembling the empirical CRISPR screens and the benchmark settings evaluated in this study is provided in Supplementary File 1. Simulation results in all figures are summarized across 25 independent simulation runs per setting.

## 5 Data & code availability

The CRISPR screening datasets analyzed in this study were obtained from previously published studies. The HDCRISPR dataset was derived from Henkel et al. (2020) (Henkel et al. 2020), with associated data and analysis resources available through the authors’ published supplementary materials and repository (https://github.com/boutroslab/Supplemental-Material/tree/master/Henkel%26Rauscher_2019). The exposure knockout screen (Doxo dataset) was obtained from Kim et al. (Kim et al. 2026), where the data are described within the publication and corresponding supplementary materials. The simulation framework and analysis code developed in this study are publicly available at: https://github.com/bachergroup/simCRISPR.

## 6 Authors’ contributions

ZZ and XD designed the `simCRISPR` methodology and software. ZZ designed and performed simulation studies, analyzed empirical CRISPR screen datasets, and wrote the manuscript. XD developed the `simCRISPR` package and assisted with manuscript writing. CK provided the empirical CRISPR screen data and assisted with biological interpretation of the case-study results.

CM contributed to the visualization of the experimental design. WBB and CDV contributed to study design and biological interpretation. RB conceived the study, contributed to the design of the `simCRISPR` simulation framework and statistical methodology, provided guidance on the computational analyses, contributed to the interpretation of results, and supervised the project. All authors contributed to manuscript writing, revising, and figure preparation. All authors discussed the results, contributed to the final manuscript, and approved its submission.

