## Supplemental File 2 for "simCRISPR: Modeling Experimental Complexity in Pooled CRISPR Screens"

### Supplementary File 2

To explore how different effect settings influence model performance, we conducted a series of simulation experiments in which one effect was varied at a time while all other parameters were held at their baseline values. The effects examined included the knockout (KO) effect, treatment effect, DNA damage effect, KO-treatment interaction effects, and DNA damage interaction effect.

For each setting, balanced accuracy (BA) for detecting interaction effects was averaged across 25 simulation runs. The following figures summarize the mean balanced accuracy under different settings for each effect.

**Figure S1. Effect of Knockout Effect on Balanced Accuracy.**
Balanced accuracy remained relatively stable across knockout effect sizes for most methods. A clearer increase was observed for **DESeq2 with SFHB normalization and thresholding**, suggesting that empirical log2FC thresholding may remove false positives, and thereby improve classification performance in this setting.

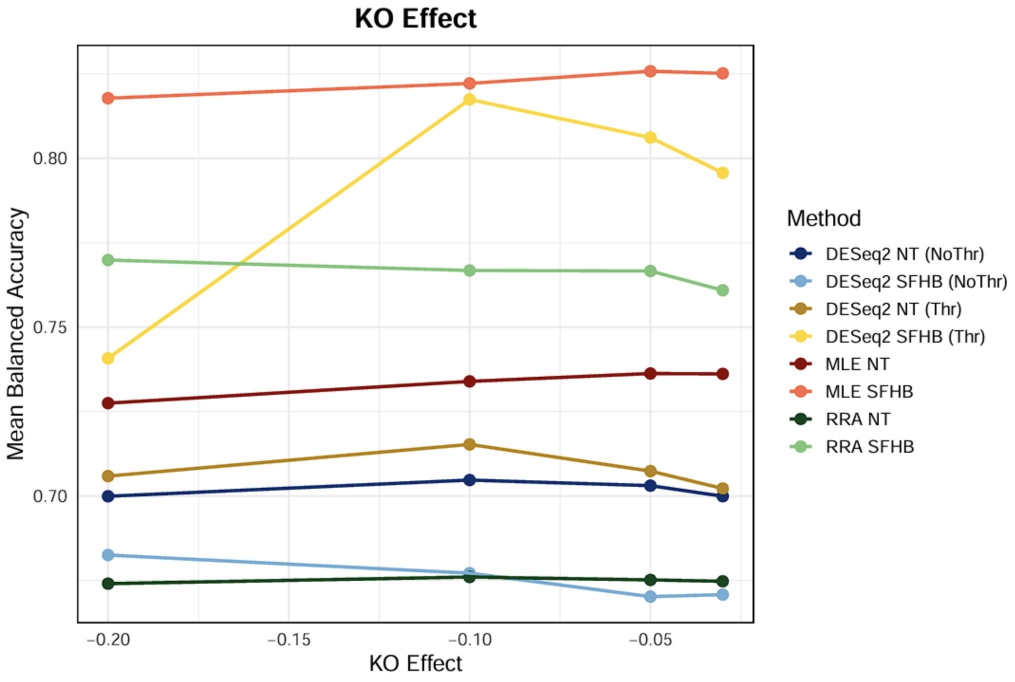

**Figure S2A. Effect of Treatment Effect on Balanced Accuracy.**
Balanced accuracy varies little across treatment effect sizes and shows no clear trend as the effect size increases.

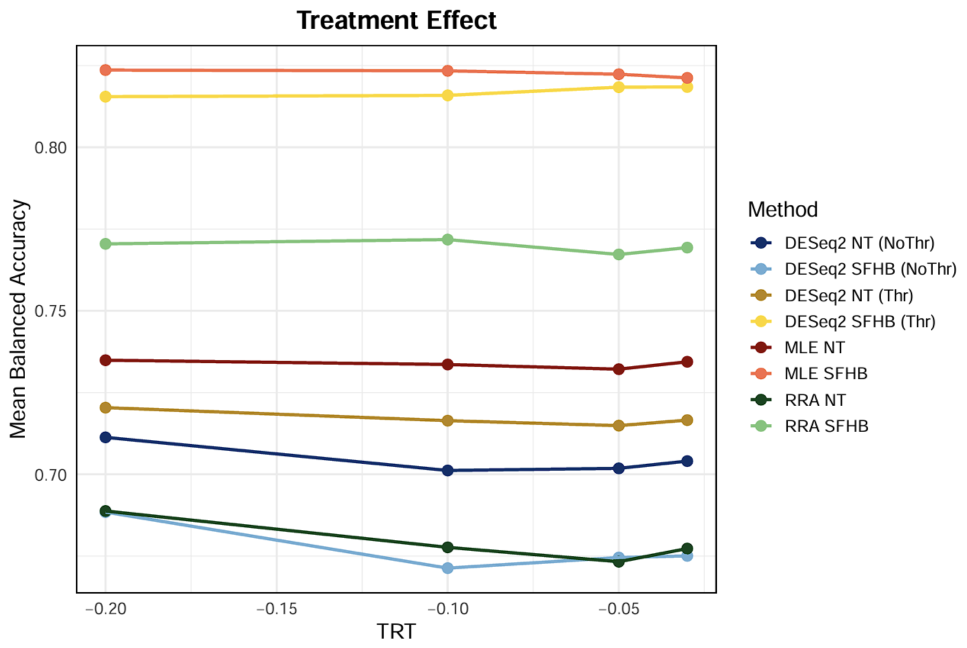


**Figure S2B. Effect of Treatment Effect Standard Deviation on Balanced Accuracy.**
Balanced accuracy decreases as the standard deviation of the treatment effect increases, indicating that higher variability makes interaction effects harder to detect.

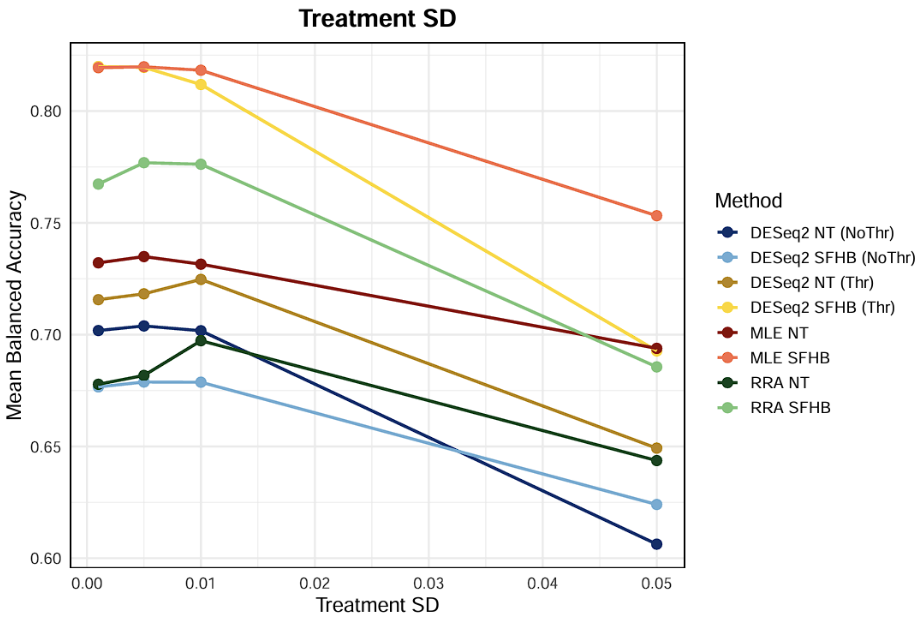

**Figure S3A. Effect of DNA Damage Effect on Balanced Accuracy.**
Balanced accuracy exhibits limited variation across DNA damage effect sizes. Similar trends are observed across settings, with only small changes as the effect size increases.

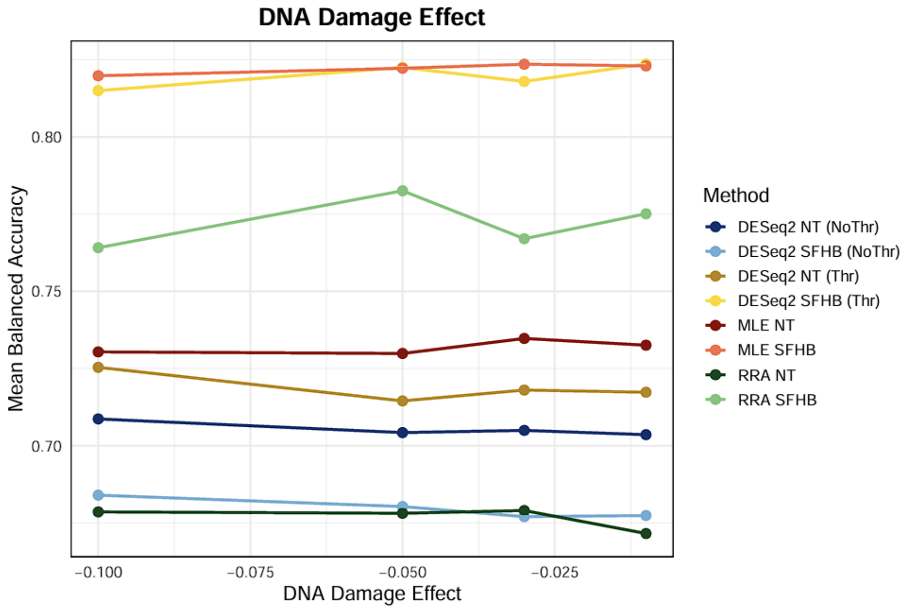


**Figure S3B. Effect of DNA Damage Effect Standard Deviation on Balanced Accuracy.**
Balanced accuracy changes only slightly as the standard deviation of the DNA damage effect increases, with similar patterns observed across methods.

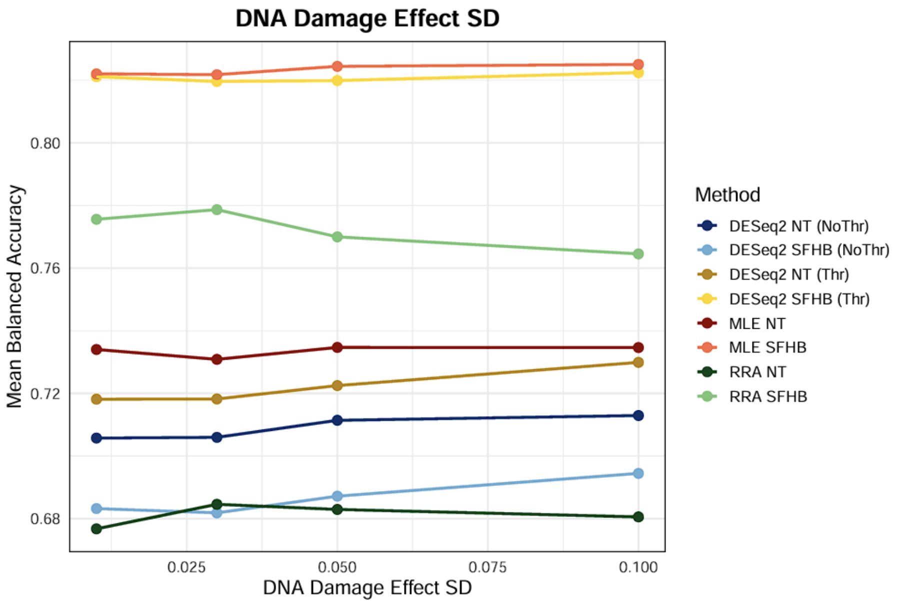

**Figure S4. Effect of KO-Treatment Interaction on Balanced Accuracy.**
Balanced accuracy increases as the interaction strength becomes stronger, reflecting improved detection of interaction effects. At higher interaction levels, DESeq2 without thresholding shows a slight decline, while applying a threshold results in more stable performance, suggesting that filtering may help reduce false positives.

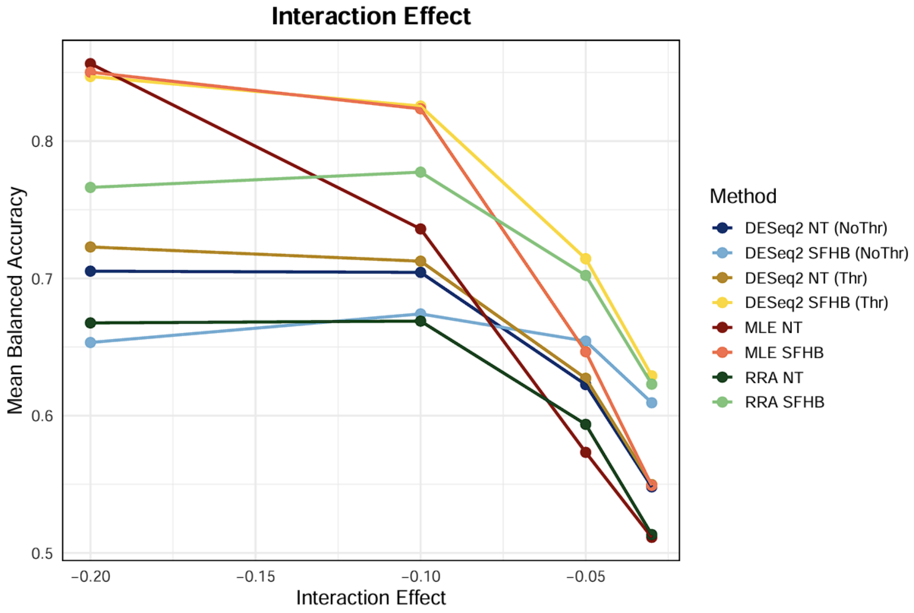


**Figure S5. Effect of DNA Damage Interaction Effect on Balanced Accuracy.**
Balanced accuracy decreases as the DNA damage interaction effect increases, with a similar downward trend observed across methods.

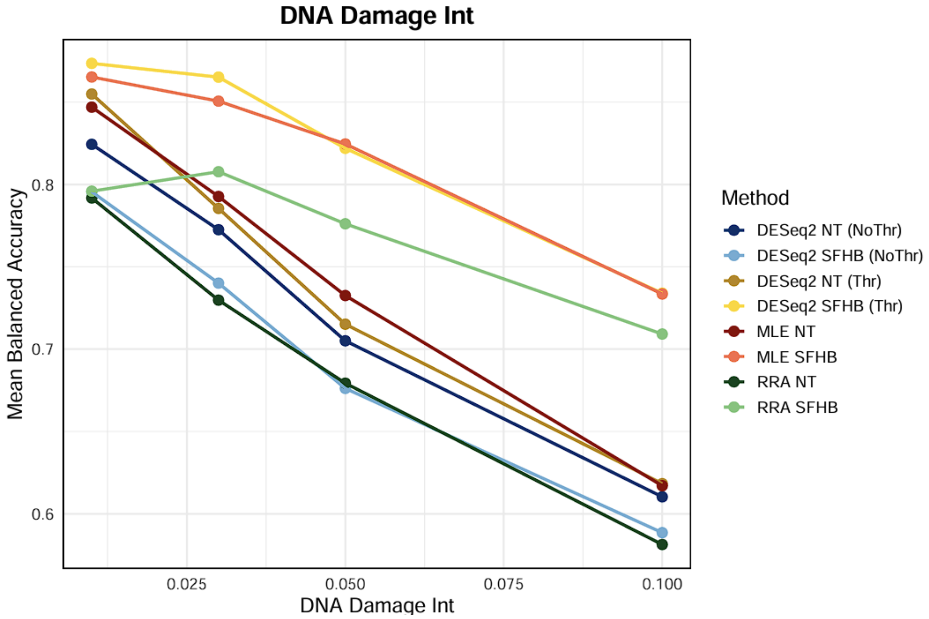
