## Supplemental File 1 for "simCRISPR: Modeling Experimental Complexity in Pooled CRISPR Screens"

### Supplementary File 1

Supplementary File 1 summarizes the simulation parameters used for the simulated datasets in Figures 2–5. Tables S1A and S1B provide the baseline parameters used to generate the representative simulated dataset shown in Figures 2C and 2F. The remaining tables describe the parameter changes used for the additional simulation analyses, including the DDR-treatment interaction strength analysis in Figure 3B and the interaction-effect benchmarking analyses in Figures 4 and 5. For each setting, 25 independent simulation runs were generated.

**Table S1A. Baseline simulation and biological-effect parameters.** These parameters define the baseline growth model, sgRNA library composition, initial sgRNA abundance, and biological effect settings used to generate the representative simulated dataset in Figures 2C and 2F. Unless otherwise specified, the later simulation analyses were generated by modifying selected parameters from this baseline setting.

| Parameter | Value | Description |
|---|---|---|
| **baseline_gr** | 0.1 | Baseline cell growth rate used in the growth model. |
| **method** | Exp (Exponential) | Growth model used for simulation cell proliferation. |
| **samples** | Independent | Experimental design for sample generation. |
| **L** | 10000 | Carrying capacity (maximum population size) used only in logistic growth model. |
| **sample_noise_SD** | 0.05 | Standard deviation (SD) of random noise across different conditions or replicates. |
| **noise_SD** | 0.01 | SD of additional technical noise. |
| **days** | 5 | Number of time points (e.g. days). |
| **reps** | 3 | Number of biological replicates per condition. |
| **n_total** | 3200 | Total number of sgRNAs. |
| **n_ntgt** | 100 | Number of non-targeting sgRNAs. |
| **n_sfhb** | 50 | Number of safe harbor sgRNAs. |

| | | |
|---|---|---|
| **initial_dist** | Normal | Distribution used to generate initial sgRNA counts at the beginning. |
| **initial_mu** | 1100 | Mean of the distribution used to generate initial sgRNA counts. |
| **initial_sd** | 300 | SD of initial sgRNA abundance. |
| **trt_eff_mu** | -0.1 | Mean of the treatment effect. |
| **trt_eff_sd** | 0.001 | SD of the treatment effect. |
| **sg_eff_mu1** | -0.1 | Mean of the negative sgRNA knockout effect. |
| **sg_eff_mu2** | 0.03 | Mean of the positive sgRNA knockout effect. |
| **sg_eff_prop1** | 0.8 | Proportion of sgRNAs with negative knockout effects. |
| **distbDNA_mu** | -0.01 | Mean of the DNA damage response effect. |
| **distbDNA_sd** | 0.01 | SD of the DNA damage response effect. |
| **distb_trt_eff_mu1** | 0.03 | Mean primary DDR-treatment interaction effect |
| **distb_trt_eff_mu2** | 0 | Mean secondary DDR-treatment interaction effect; not used when distb_trt_eff_prop1 = 1 |
| **distb_trt_eff_prop1** | 1 | Proportion assigned to the primary DDR-treatment interaction component. |
| **sg_trt_eff_mu1** | -0.1 | Mean negative KO-treatment interaction effect |
| **sg_trt_eff_mu2** | 0.1 | Mean positive KO-treatment interaction effect |
| **sg_trt_eff_prop1** | 0.8 | Proportion of KO-treatment interaction effects that are negative |

**Table S1B. Baseline PCR amplification and sequencing parameters.** These parameters define the PCR amplification, sequencing-depth, and library equalization settings applied consistently across all simulation analyses.

| Parameter | Value | Description |
|---|---|---|
| **rounds** | 5 | Number of PCR amplification rounds to simulate. |
| **totalDepth** | $2 \times 10^7$ | Total sequencing depth to distribution across sgRNAs. |
| **cm_mu** | 0.99 | Mean of capture efficiency. |
| **cm_sd** | 0.01 | SD of the capture efficiency. |
| **ep_mu** | 0.01 | Mean of the error propagation rate during PCR. |
| **ep_sd** | 0.001 | SD of the error propagation rate. |
| **pcr_sd** | 0.02 | SD of PCR amplification noise. |
| **sf_sd** | 0.2 | SD for sampling fluctuation. |

**Table S2. DNA damage-treatment interaction strength settings used for Figure 3B.** For Figure 3B, only *distb_trt_eff_mu1* was varied; all other parameters were kept at the baseline settings in Tables S1A and S1B.

| Parameter | Values tested |
|---|---|
| **distb_trt_eff_mu1** | -0.20, -0.10, -0.07, -0.03, 0, 0.03, 0.07, 0.10, 0.15, 0.20 |

**Table S3. GxE interaction effect settings without DNA damage-related effects for Figure 4.** For Figure 4, DNA damage response and DDR-treatment interaction effects were excluded. KO-treatment interaction strength was varied by changing *sg_trt_eff_mu1*, while all other parameters were kept at the baseline settings in Tables S1A and S1B.

| Parameter | Values tested |
|---|---|
| **sg_trt_eff_mu1** | -0.05, -0.10, -0.20 |

**Table S4. GxE interaction effect settings with DNA damage-related effects for Figure 5.** For Figure 5, DNA damage response and DDR-treatment interaction effects were included at fixed values. KO-treatment interaction strength was varied by changing *sg_trt_eff_mu1*, while all other parameters were kept at the baseline settings in Tables S1A and S1B.

| **Parameter** | **Value(s) tested** |
|---|---|
| **sg_trt_eff_mu1** | -0.05, -0.1, -0.2 |
| **distbDNA_mu** | -0.01 |
| **distbDNA_sd** | 0.01 |
| **distb_trt_eff_mu1** | 0.05 |
